# Gamma Oscillations and Emotional Memory

**DOI:** 10.1101/2022.02.02.478869

**Authors:** Lisa Luther, Lycia D. de Voogd, Muriel A. Hagenaars, Ole Jensen

## Abstract

Increased visual gamma band activity is known to relate to memory and encoding in cognitive tasks and has been shown to be present during arousing compared to neutral stimulus processing. Here, we set out to investigate whether gamma activity is stronger for later remembered compared to later forgotten emotional pictures. Thirty-two healthy participants underwent a passive viewing task in which they viewed 208 (104 arousing (52 pleasant and 52 unpleasant) and 104 neutral) pictures from the International Affective Picture System (IAPS) database while we simultaneously recorded their electroencephalogram (EEG). Immediately after, they performed a recognition task including the 208 target pictures they saw before and 104 new pictures (52 arousing (26 pleasant and 26 unpleasant) and 52 neutral) as lures. We found that arousing pictures (unpleasant ones in particular) were better remembered compared to neutral pictures. Importantly, gamma activity was enhanced for remembered compared to forgotten pictures and this effect was stronger for arousing pictures. We conclude that previous findings regarding enhanced gamma as a signature of subsequent memory generalizes to emotional memory. Our findings therefore shed new light on the role of visual gamma band activity in arousal and memory formation.

## Introduction

Memories for arousing events are typically well-remembered (Bradley et al., 1992; Cahill & McGaugh, 1996; Hermans et al., 2011; Schacter, 1999). In the first place, this selectivity of memory can be explained by the immediate effects of arousal on attentional, sensory, and subcortical mnemonic processes (Bradley et al., 1992; Davis & Whalen, 2001). On a cortical level, gamma activity is known to reflect increased attention and neuronal information processing (Fries et al., 2001; Tallon-Baudry & Bertrand, 1999) and increased memory encoding (Gruber et al., 2004; Osipova et al., 2006; Pulvermüller et al., 1999). Gamma activity is also increased during the perception of arousing events compared to neutral events, independent of memory (Aftanas et al., 2004; Balconi et al., 2009; Keil et al., 2001). It is unknown, however, whether increases in gamma activity support the increased memory formation found for arousing events.

Differential neural activity between later remembered versus later forgotten information is referred to as the subsequent memory effect (for review see Paller & Wagner, 2002). Increased gamma power during (incidental) encoding has been shown to predict whether the picture will later be remembered or forgotten in a wide array of non-emotional cognitive tasks. Osipova and colleagues (2006) found increased gamma for landscapes and buildings when they were later remembered compared to later forgotten. Gruber et al. (2004) found an increased gamma power for later remembered over forgotten words. Similarly, another study using intracranial recordings, found an increase in gamma activity widely spread over the cortex to predict later remembered compared later forgotten words (Sederberg et al., 2003). For a review on oscillatory brain processes during subsequent memory formations see Hanslmayr and Staudigl (2014). Thus, gamma activity is suggested to underlie increased attention during encoding that strengthens memory.

Gamma activity also plays a role in affective attention. Previously, we found a valence-specific effect for gamma: gamma activity was enhanced during the perception of unpleasant compared to neutral and pleasant pictures (Luther et al., in prep.). Similarly, Oya and colleagues (Oya et al., 2002) found gamma activity to be enhanced for unpleasant pictures (IAPS) but not for neutral or pleasant pictures compared to baseline when analyzing data from depth electrodes in the human amygdala. However, Keil et al. (2001) found a valence-independent effect for arousal: increased gamma activity (45 – 60Hz) over right posterior sensors for arousing (pleasant and unpleasant) compared to neutral pictures. Also, Mueller, Keil, Gruber, and Elbert (1999) reported increased gamma power (30 – 50 Hz) over right frontal and temporal sites for arousing (both pleasant and unpleasant) compared to neutral pictures, but no effect was observed for gamma power between 50 – 90 Hz. Greater gamma band power was also observed during emotional (fearful) relative to neutral faces (Luo et al., 2009). Thus, gamma activity seems to be associated with arousal, and possibly also with valence. We therefore hypothesized that gamma activity underlies the enhanced memory found for arousing events. Due to limitations in the study design, we cannot test a difference between valence and arousal.

To describe the role of gamma activity in emotional memory we tested whether later remembered arousing pictures are characterized by increased gamma activity at encoding. We conducted an EEG study using pictures from the IAPS data base (neutral and arousing (unpleasant and pleasant)). 208 Pictures were viewed while the EEG was measured. This was followed by a recognition task including all pictures and an additional 104 new pictures as lures. We analyzed gamma power during the encoding phase in a 2 by 2 design: later memory outcome (later remembered; LR / later forgotten; LF) by arousal (arousing / neutral).

First, replicating earlier studies, we predicted increased gamma activity for pictures later remembered compared to pictures later forgotten. Second, also replicating earlier studies, we predicted increased gamma activity for arousing (pleasant and unpleasant) compared to neutral pictures. Lastly, we predicted an interaction such that the increase in gamma activity for pictures later remembered would be stronger for the arousing pictures.

## Methods

### Participants

Fourty-four participants (18 male) were recruited from Radboud University Nijmegen, via an experiment participation system, flyers, and personal contacts. Due to technical problems leading to incomplete data sets, data from four participants were excluded. Additionally, three more participants dropped out during the course of the experiment due to dizziness or feeling unwell, and 7 datasets contained too few trials without EEG artifacts (< 10 trials in one cell). The mean age of the 30 remaining participants was 23.6 (*SD = 8.5*). The study was approved by the local ethics committee (Ethische Commissie Gedragswetenschappelijk onderzoek, ECG) and participants gave written informed consent prior to the experiment.

### Data Acquisition

#### Apparatus

During encoding, participants were standing while the EEG was recorded. Stimuli were presented on a Samsung 2233SW screen with a refresh-rate of 60Hz or 120Hz, which was adjusted to eye-height. A 64 channel EEG system (ActiCap64 system, Brain Products GmbH, Gilching, Germany) based on the extended 10-10 system was applied. The signals were referenced to linked mastoids and later re-referenced to a common reference. The data were sampled at 500 Hz, following band-pass filtering between 0.016 and 150 Hz.

### Procedure

Participants took part in a larger study, of which the third task was a passive viewing task containing International Affective Picture System pictures (IAPS; Lang, Bradley, & Cuthbert, 2001). The fourth task was a recognition memory task for the pictures of the passive viewing task, the data of which are presented here.

#### Stimuli

During the passive viewing task each participant was presented with 208 IAPS pictures of which 52 were pleasant, 52 unpleasant, and 104 neutral pictures. During the recognition task that followed, these 208 were presented again, mixed with 104 lures, leading to 312 pictures in total (78 per emotion and 152 neutrals, which included the targets and 50% lures). Those 312 pictures constitute the entire picture set of the study; whether a given picture was used as a target or lure for a given participant was determined via a rotation list balanced across subjects. Pictures were selected based on the normative valence ratings provided by the IAPS manual (cut-off values: unpleasant < 3.2; neutral: 4 – 6; pleasant > 6.8). The valence ratings were further constrained not to differ more than 0.8 points between men’s and women’s ratings. The pictures were matched in terms of luminance and size with a custom written program. The same amount of pictures including humans (including the face) and non-humans was included in each category (double the amount in the neutral category, respectively).

#### Passive viewing task

Each picture appeared only once during the passive viewing task. Pictures were presented in 8 blocks consisting of 26 pictures. Each block contained 13 neutral pictures and 13 emotional pictures. This amounted to 4 blocks of pleasant and neutral pictures, and 4 blocks of unpleasant and neutral pictures. Thus, each block consisted half of neutral and half of arousing pictures (of either valence). The pictures and blocks were pseudo-randomized, being constrained to not show more than 2 blocks of one valence (pleasant or unpleasant) sequentially, and within each block not more than 4 pictures of one valence (neutral or pleasant, or neutral or unpleasant, respectively) in a row. Pictures were presented for 2 s with inter-trial intervals (ITI) of 1.5 – 2 s, leading to a total duration of around 1 min 36 s per block. A fixation-cross stayed on the screen and the pictures were presented around it (Figure 1). The blocks were separated by 20 s breaks. After 4 blocks followed a 3 min break allowing participants to sit down and rest their legs. During the break this cross turned into a countdown before resuming the picture presentation. Participants were instructed to fixate on the cross while perceiving the picture around it. Furthermore, they were instructed to blink after each picture in order to reduce blinking during stimulus presentation. Their posture was required to be relaxed with arms hanging alongside the body.

**Figure 1.**
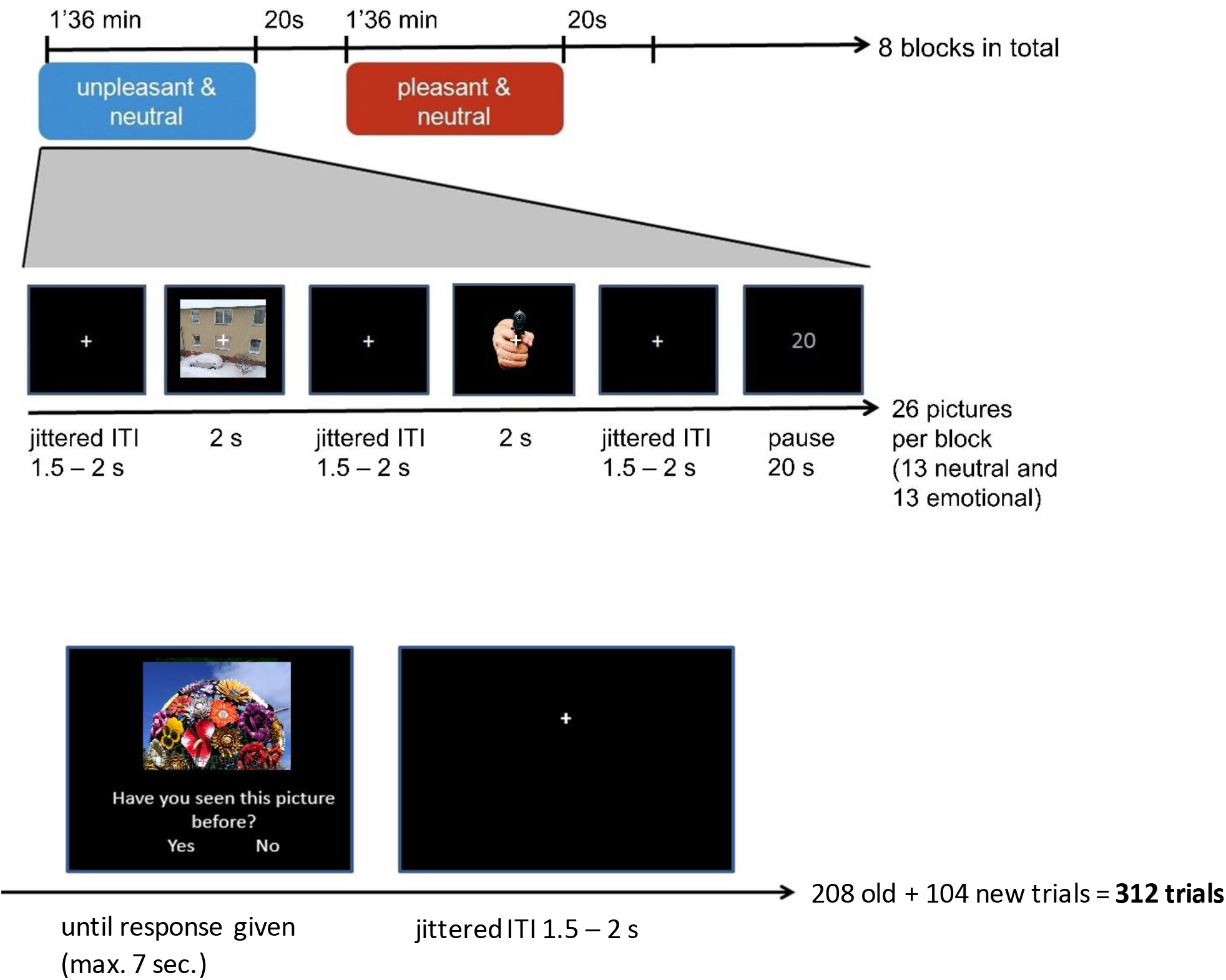
Experimental paradigm. Upper panel: Passive viewing task during which participants were gazing at the fixation cross while perceiving the picture around the cross. Neutral pictures were presented among either pleasant or unpleasant pictures. Lower panel: Recognition memory was assessed after all IAPS pictures had been viewed.

#### Picture recognition memory task

The recognition paradigm contained all 208 pictures presented during the passive viewing paradigm (targets) with an additional 104 new pictures (lures). Pictures were presented until response was given, but no longer than 7 s. Of the 104 lures pictures, 26 were unpleasant, 26 were pleasant and 52 were neutral. Participants were instructed to indicate whether they had seen the picture before, or whether it was a new picture and responded by pressing a button with their right hand index finger for ‘yes’ and with their middle finger for ‘no’. Pictures were presented in a consecutive order in 8 blocks of 39 pictures with 20-second breaks in between and a 3 minutes’ break after 156 pictures. The presentation order of targets and lures was fully random. Since 2/3 of the pictures were ‘old’ and 1/3 was ‘new’, chance level of responding to old pictures as ‘already seen’ was at 66%.

### Data Analysis

#### Subsequent memory analysis

For the subsequent memory analysis, each of the 208 pictures were assigned to whether they were later remembered or later forgotten. Memory accuracy was assessed by subtracting the hit rates (i.e., ‘old’ response to a target) from the false alarm rates (i.e., ‘old’ response to a lure) for each condition. For the EEG analysis, we collapsed both unpleasant and pleasant valance together, due to a high memory performance for each valence separately, leading to too few forgotten pictures for EEG analysis.

#### EEG sensor level analyses

EEG data were analyzed using MatLab (MathWorks Inc., 2014b) and the Matlab-based FieldTrip toolbox (Oostenveld, Fries, Maris, & Schoffelen, 2011). Artifacts were removed in a semi-automatic manner using visual inspection and selecting a threshold for each participant (Horschig et al., 2015). Trials containing ocular or muscle-artifacts were detected based on this threshold and excluded from analysis (on average 31%). Datasets with too many lost trials (less than 10 trials left in one cell) were excluded from the EEG analysis (7 participants; also see participants section).

Sensor level time-frequency analysis was performed using a 400 ms sliding time window and a multi-taper approach (discrete prolate spheroidal sequences; DPSS, Percival & Walden, 1993) for the frequencies from 30 – 120 Hz (Osipova et al., 2006).

#### EEG statistical analyses

Statistical analysis was performed on the whole brain sensor level. For each participant, trials were averaged depending on the response given during the recognition session (LR/LF) or by condition (arousing / neutral) or by both (2×2: LR/LF x arousing/neutral). To assess whether sensor level power is significantly different between later remembered (LR) and later forgotten (LF) trials, a cluster based non-parametric permutation test was used within subjects (Maris & Oostenveld, 2007). This test clusters neighboring sensors and time-frequency points which show the same effect. For each grid point a dependent samples *t*-test (LR vs. LF) was computed. For each sample, the *t*-value is calculated. Clusters are formed based on spatial adjacency of grid points where values exceed the critical threshold of *p* = 0.05 (uncorrected). The sum of the *t*-values within a cluster was used as cluster-level statistic and the cluster with the maximum sum was used for the test statistic. To create a reference distribution 1000 permutations of randomization were performed (van Dijk et al., 2008). By conducting only one comparison (between the actual cluster size and the distribution of cluster sizes achieved through the permutations) it controls for multiple comparisons. The test was conducted over the full window of - 1000ms to 2000ms around stimulus onset, applying a baseline from the −800 to −200ms interval, allowing for data-driven discovery of differences in terms of channels, time points, and frequencies.

## Results

### Subsequent memory: behaviour

Memory accuracy in the picture recognition memory paradigm was assessed by subtracting the hit rates from the false alarm rates for the unpleasant, pleasant, and neutral pictures. Overall performance was above chance level (overall hit rate versus false alarm rate; F(1,36) = 876.10, p < .001, Pη2 = .96). There was an accuracy difference between the three valence categories (F(1,36) = 52.19, p < 001, Pη2 =.59). Unpleasant pictures lead to higher memory accuracy than pleasant (t(36) = 7.823, p < .001) and neutral (t(36) = 7.924, p < .001) pictures. There was no difference in memory accuracy between the pleasant and the neutral pictures (t(36) = 1.185, p = .24). In a separate t-test, arousing pictures (pleasant an unpleasant combined) were better remembered than neutral pictures (t(36) = 6.22, p < 0.001). See Figure 2 and Table 1 for descriptive statistics.

**TABLE 1.** Memory accuracy: number of trials (subsequent memory performance per category).

|  | Remembered |  | Forgotten |  |
| --- | --- | --- | --- | --- |
|  | Mean | SD | Mean | SD |
| Unpleasant | 42.38 | 4.84 | 9.62 | 4.84 |
| Pleasant | 32.92 | 8.86 | 19.08 | 8.86 |
| *Total arousing* | 75.30 | 12.36 | 28.70 | 12.36 |
| Neutral | 63.70 | 17.63 | 40.30 | 17.63 |

**Figure 2.**
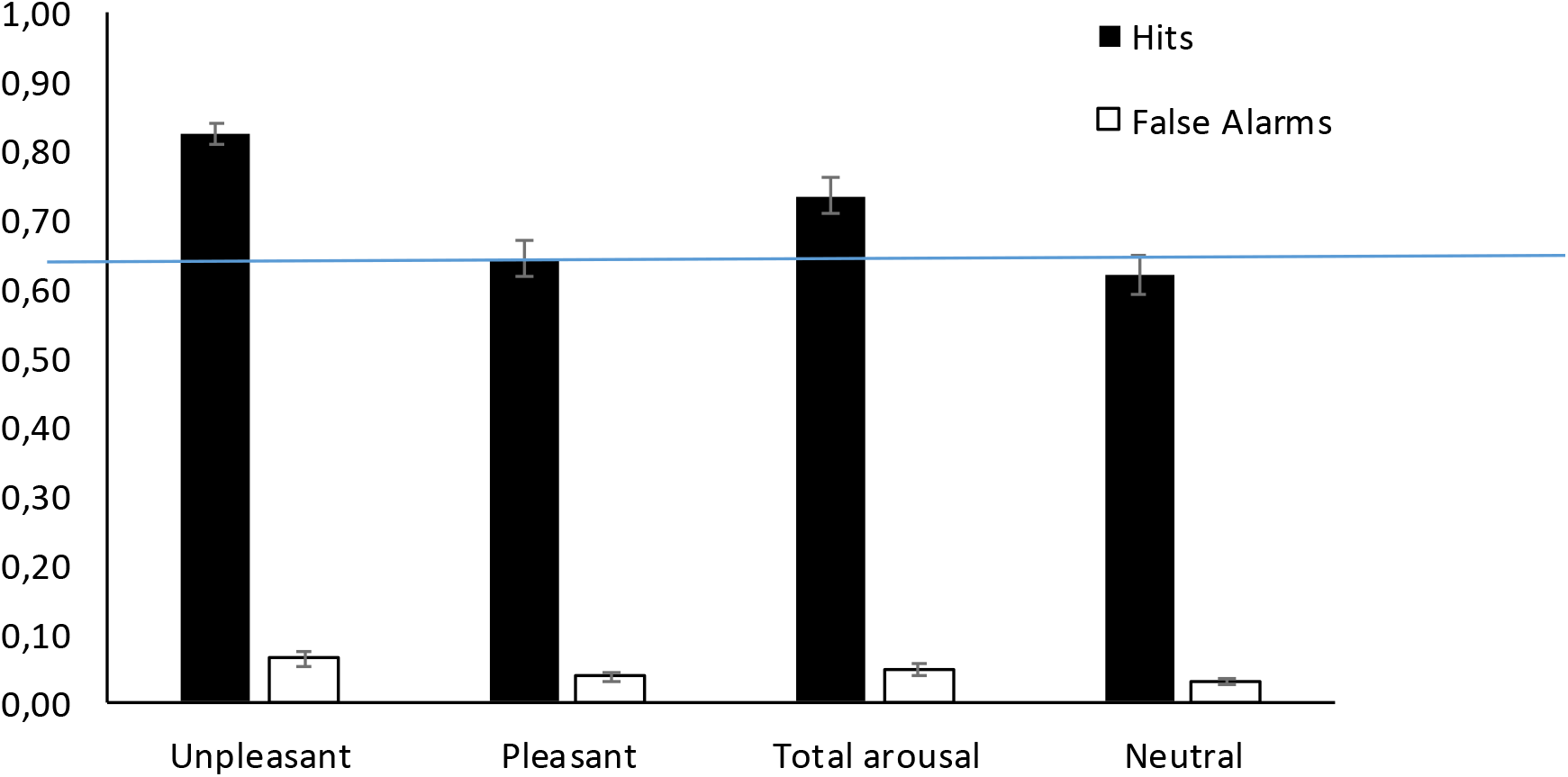
Memory performance. Overall memory performance (hit rate versus false alarm rate) is above chance level (indicated by the blue line). Arousing pictures (unpleasant and pleasant combined) were remembered significantly better than neutral ones. Error bars indicate standard error of the mean (SEM).

### EEG

#### Stronger gamma for remembered compared to forgotten pictures

Gamma power differences related to the subsequent memory effect were identified by comparing later remembered with later forgotten pictures (independent of picture category; Figure 3) in a cluster-based permutation test allowing for bottom-up discovery of significant time points, frequencies, and sensors. We observed stronger gamma band activity for LR compared to LF over central posterior areas (p < .001). This was observed in posterior sensors in the 500 – 2000ms interval in the 50 – 80 Hz and 100 – 110 Hz frequency range. Thus, we were able to replicate previous findings of enhanced gamma during encoding for successfully remembered items.

**Figure 3.**
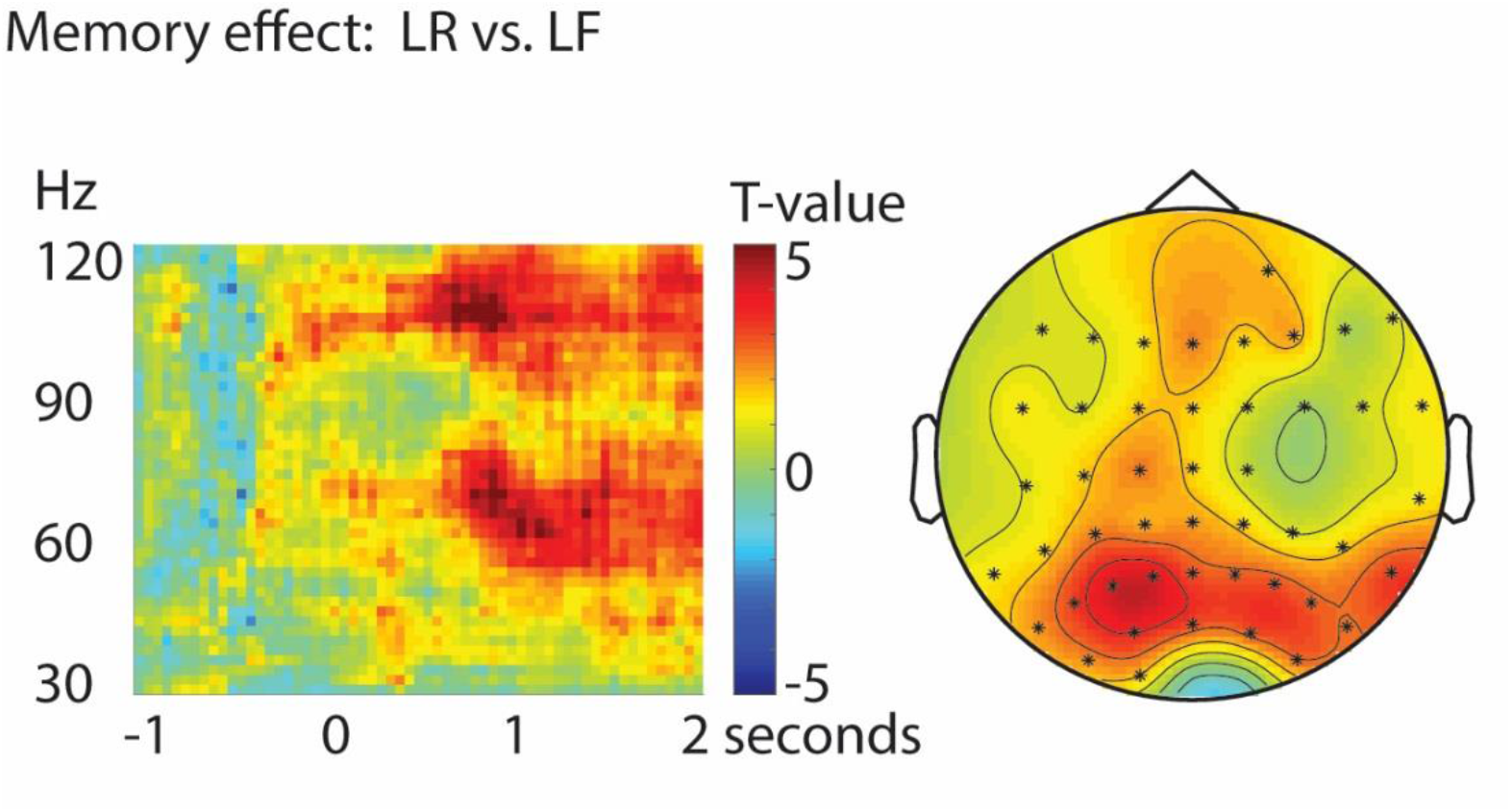
Time-frequency representation (TFR) and topographical representation of statistical test. The TFR is shown for electrode Pz. Sensors for which there was a significant difference in the named contrast are indicated with a dot in the topographic plot: nearly all channels reached significance; the effect being most pronounced in the 500 – 2000ms time window between 50 – 80 and 100 – 110 Hz. Timepoint 0 = picture onset time.

#### Increased gamma for arousing compared to neutral pictures

Gamma power differences related to arousal were identified by comparing arousing (unpleasant and pleasant) with neutral pictures, independent of later memory outcome. We observed stronger gamma band activity for arousing compared to neutral pictures (p < .001; see Figure 4). This effect was most pronounced over posterior sensors in the 0 – 2000ms interval and the 70 – 80 Hz frequency range. Thus, valence independent arousing pictures yield more gamma activity during encoding.

**Figure 4.**
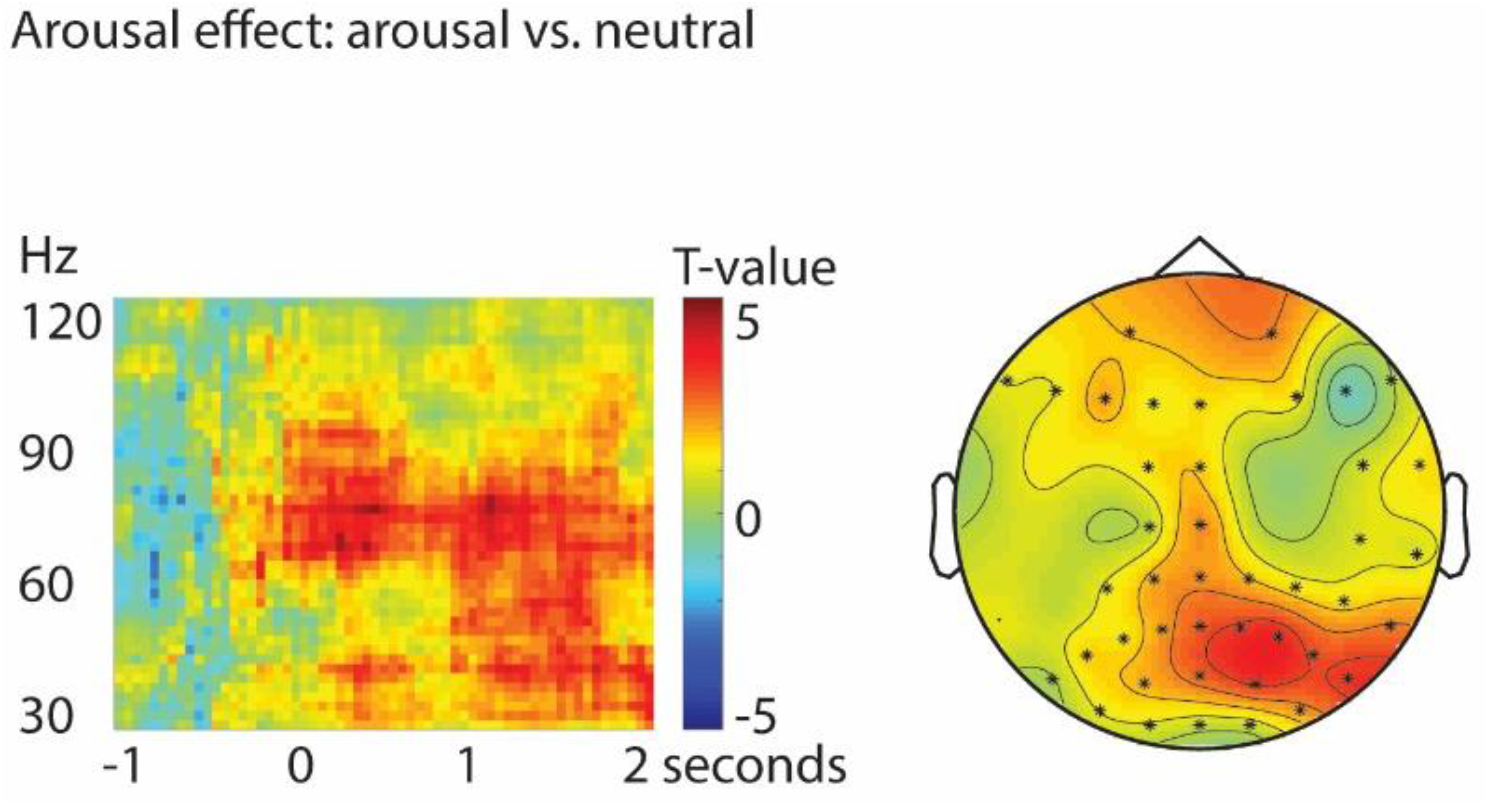
Time-frequency representation (TFR) and topographical representation of statistical test. The TFR is shown for electrode Pz. Sensors for which there was a significant difference in the named contrast are indicated with a dot in the topographic plot.

#### Gamma for remembered pictures stronger for the arousing pictures

The interaction effect between emotion and memory showed a statistical trend at p = .060 (see Figure 5). In line with our research question we tested the contrast of LR versus LF for the arousing pictures. We observed stronger gamma band activity for LR compared to LF pictures (p < .01). This effect was mainly pronounced over posterior sensors. For the LR pictures, arousing pictures lead to higher gamma activation than neutral stimuli (p < .01). The other two contrasts (LR versus LF in neutral and arousing versus neutral in LF) were not significant (p = .29 and p = .66, respectively). Thus, arousal indeed seems to be associated with an increase in gamma band activity in the case that the picture is later remembered, but not when it is not remembered. In the Discussion section we will go into more detail on the possible reasons for this. Also, the association of gamma with LR versus LF pictures seems to hold only for arousing pictures, while it seems that neutral pictures do not lead to a significant difference in gamma power modulation depending on memory outcome.

**Figure 5.**
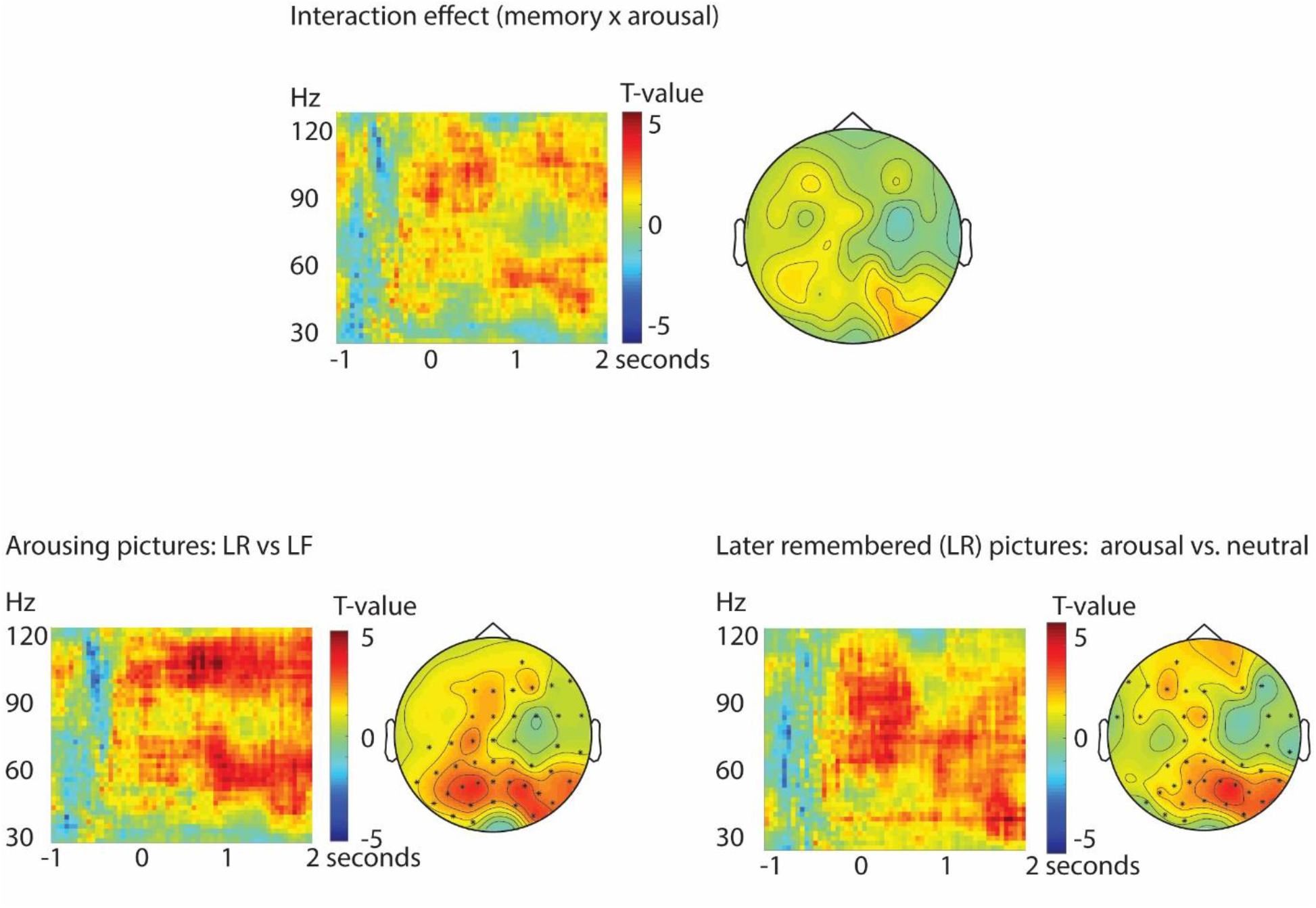
Time-frequency representations (TFRs) and topographical representations of statistical test. The TFRs are shown for electrode Pz. Sensors for which there was a significant difference in the named contrast are indicated with a dot in the topographic plot

Additionally, since other studies find effects of theta and alpha power during encoding predicting subsequent memory effects (for review see Hanslmayr & Staudigl, 2014), we ran the same tests on the lower frequencies (1 – 30 Hz; using Hanning taper instead of DPSS for the Fourier transform), but did not find an effect there (all p-values > .05).

## Discussion

We aimed to test whether gamma predicts enhanced memory for arousing stimuli. We replicated the effect of increased gamma activity for pictures later remembered compared to later forgotten, and of increased gamma activity for arousing pictures (pleasant and unpleasant) compared to neutral pictures. We also found increased gamma within remembered pictures for the arousing pictures, but not for the neutral pictures. In short, our data show that the effect of enhanced gamma reflecting subsequent memory holds for arousing pictures as well.

Confirming our first hypothesis, we found gamma activity – in its presumed role to reflect active information processing – to be enhanced during encoding, which is replicating the effect of enhanced gamma activity in relation to later remembered pictures (Gruber et al., 2004; Hanslmayr & Staudigl, 2014; Osipova et al., 2006; Sederberg et al., 2003). This still holds when looking at arousing pictures separately.

Confirming our second hypothesis, we found gamma to be enhanced towards arousing compared to neutral pictures, which is in line with literature on affective processing (Keil et al., 2001; Luo et al., 2009; Mueller et al., 1999). This consolidates findings showing that arousing items draw attention and the above mentioned role of gamma activity reflecting active processing in the cortex.

Interestingly, we showed an interaction effect between arousal and memory at trend level (p = .060) and the planned contrasts showed significantly enhanced gamma activation both for remembered versus forgotten arousing pictures, as well as arousing versus neutral remembered pictures.

### Later forgotten pictures

The contrast of the forgotten pictures (arousing vs. neutral) did not show a significant effect. This contrast is relatively noisy, because 1) the signal-to-noise ratio of the data is noisier than in the other cells due to the smaller amount of trials, and 2) it is unclear what the participants were doing at the moment of seeing a picture which they did not encode in memory – they may not have been attentive to the picture, but we inherently cannot know what they were processing instead.

### Neutral pictures

The subsequent memory effect that we observe in the grand average for arousing pictures is not apparent when comparing only the neutral pictures with each other (LR vs. LF). The absence of a difference in gamma activity for LR vs. LF pictures of the neutral condition could be a result of the semi-blocked design. Neutral pictures are remembered less well when surrounded by emotional pictures (Watts et al., 2014). Watts and colleagues investigated the difference between subsequent memory effects of emotional pictures presented in blocked (unpleasant/neutral) and intermixed (unpleasant and neutral pictures in one block) presentation of pictures. Later remembered pictures elicited a larger ERP (event-related potential) as of 300ms after stimulus onset. This memory effect still holds for emotional and neutral pictures independently, and for pictures presented in pure blocks and in mixed blocks, except for neutral pictures in mixed blocks over posterior sites, indicating that emotional pictures are remembered equally well independent of the surrounding pictures, whereas neutral pictures are remembered less well when surrounded by emotional pictures. In our study we used a mixed design, which according to the findings of Watts and colleagues induces a memory decrease of neutral information. This could have been part of why we found less gamma for the neutral pictures.

### A central posterior effect

It is probably not surprising that we found our effect to be strongest over the posterior areas, since our task was visual. Lakatos et al. (2004), found attention to be reflected strongest over the brain area which’s modality was being used, which in our case is the visual domain, being processed mainly in occipital and adjacent regions.

### Future directions

Our study used purely visual input to reveal that enhanced gamma activity is at its largest for arousing, later remembered items. From this, the testable prediction arises that this might not only be true in the visual but also in the auditory domain, and for linguistic input (written or spoken sentences). Future work might aim to replicate and solidify the findings in this manner. Finally, it would be good to investigate the oscillatory effect per valence (unpleasant/pleasant), especially since our data show that unpleasant pictures are better remembered. Given that particularly unpleasant trials are best remembered and that arousing, later remembered trials elicit most gamma power, this research could be used in the field of trauma memory development. Potentially, gamma power difference scores in combination with memory performance difference scores between unpleasant, neutral and pleasant input, could be predictive of later post-traumatic stress disorder development. A cohort study among a demographic likely to be exposed to traumatic events (e.g. military personnel) could shed light on this.

### Mechanistic underpinnings of our effects

Memory for arousing, unpleasant events has been strongly linked to amygdala activation (Hermans et al., 2014). Gamma activity has also been recorded in the amygdala during fearful events (Oya et al., 2002), but phyisologically these deep signals cannot be picked up by scalp electrodes such as the ones we employed. Thus, the question arises whether, and if so, how, deep-lying structures such as the amygdala could be linked to our gamma findings. Possibly, when the amygdala interacts with the cortex (Hermans et al., 2014; Roy et al., 2009), it promotes processing of the arousing pictures over the neutral ones, contributing to the enhanced gamma we find for the arousing pictures. In their review paper, Tamietto and DeGelder (2010) describe cortical and subcortical pathways for vision and emotion. Apart from the primary visual pathway via the occipital lobe, visual information with threatening content takes a route via the amygdala which in turn projects via several midbrain nuceli (namely the substantia nigra, nucleus basalis of Meynert, and the locus coeruleus) onto large parts of the neocortex. Early visual information also enters the superior colliculus, which projects onto these midbrain nuclei and the pulvinar, which projects to frontoparietal areas. Both pathwaysmay underlie the enhanced activity in the gamma band which we observe for arousing, later remembered items.

### Limitations

A limitation of the present study lies in the fact, that in our design there were too few forgotten emotional pictures for both arousing valences, and hence a valence-based analysis of the EEG was not possible. Since in our study on neural correlates during passive viewing (the encoding phase) we found gamma power to be enhanced for unpleasant compared to pleasant pictures. Likewise, in the present study, there was a valence effect on memory performance: unpleasant pictures were better remembered than neutral and pleasant pictures. Given the finding of our previous study and the current behavioural finding, it would be interesting to design a future study such that valence and arousal can be differentiated in the EEG.

In conclusion, our findings corroborate the interpretation that gamma indeed plays a role in attention-enhanced memory formation for arousing stimuli. This poses an expansion of the current knowledge base, showing for the first time that enhanced gamma for remembered items also applies to arousing stimuli.

